# Behavioral and neural correlates of diverse conditioned fear responses in male and female rats

**DOI:** 10.1101/2024.08.20.608817

**Authors:** Julia R. Mitchell, Lindsay Vincelette, Samantha Tuberman, Vivika Sheppard, Emmett Bergeron, Roberto Calitri, Rose Clark, Caitlyn Cody, Akshara Kannan, Jack Keith, Abigail Parakoyi, MaryClare Pikus, Victoria Vance, Leena Ziane, Heather Brenhouse, Mikaela A. Laine, Rebecca M. Shansky

**Author notes:** **Corresponding Author:** Rebecca M. Shansky, **Address:** 360 Huntington Ave, Boston, MA 02115.

## Abstract

Pavlovian fear conditioning is a widely used tool that models associative learning in rodents. For decades the field has used predominantly male rodents and focused on a sole conditioned fear response: freezing. However, recent work from our lab and others has identified darting as a female-biased conditioned response, characterized by an escape-like movement across a fear conditioning chamber. It is also accompanied by a behavioral phenotype: Darters reliably show decreased freezing compared to Non-darters and males and reach higher velocities in response to the foot shock (“shock response”). However, the relationship between shock response and conditioned darting is not known. This study investigated if this link is due to differences in general processing of aversive stimuli between Darters, Non-darters and males. Across a variety of modalities, including corticosterone measures, the acoustic startle test, and sensitivity to thermal pain, Darters were found not to be more reactive or sensitive to aversive stimuli, and, in some cases, they appear less reactive to Non-darters and males. Analyses of cFos activity in regions involved in pain and fear processing following fear conditioning identified discrete patterns of expression among Darters, Non-darters, and males exposed to low and high intensity foot shocks. The results from these studies further our understanding of the differences between Darters, Non-darters and males and highlight the importance of studying individual differences in fear conditioning as indicators of fear state.

## 1. Introduction

Pavlovian fear conditioning is a commonly used paradigm that models associative learning in rodents. Freezing, the total lack of all movement except that required by respiration (Fanselow, 1984), is the most common conditioned response (CR) used to indicate fear learning. The reliance on a single response as the sole indicator of learned fear reduces the likelihood of capturing and understanding the contribution of individual differences to fear learning outcomes. For example, darting is a female-biased CR that is characterized by a quick, escape-like movement across the fear conditioning chamber in response to the conditioned stimulus (CS) (Gruene et al., 2015). Our lab has found that females in general, and Darters in particular, show heightened unconditioned responses, such as shock response (how quickly the animal moves in response to the shock) to the unconditioned stimulus (US) (Gruene et al., 2015; Mitchell et al., 2022). This heightened shock response predictably emerges before darting itself does (Mitchell et al., 2022), suggesting that sex differences in shock processing may underlie the tendency of females to engage in conditioned darting.

As the recognition of darting as a conditioned response has grown, skeptics have questioned whether Darters are simply more innately stressed or hyperreactive than Non-darters and males. If so, this increased stress or hyperreactivity could be the cause of Darters’ heightened shock response, and darting could simply be one reflection of sex differences in aversive stimuli processing, which have been shown to exist across a large range of behavioral assays (Borkar et al., 2020; Greiner et al., 2019; Johnston & File, 1991; Kokras & Dalla, 2014; Laine et al., 2022; Lopez-Aumatell et al., 2008; Toufexis et al., 2016; Zambetti et al., 2019). The relationships between behavior following aversive stimuli exposure and conditioned darting have not been studied.

Sex differences in conditioned and unconditioned responses during Pavlovian fear conditioning could be a result of differences in the processing of pain from the foot shock. Unfortunately, most preclinical studies of pain sensitivity and reactivity have focused on behaviors exhibited by male rodents (Gregus et al., 2021; Mogil & Chanda, 2005), even though there are known differences in pain processing across the sexes (Bartley & Fillingim, 2013; Berkley, 1997; Blanton et al., 2021; Fillingim, 2003; Fullerton et al., 2018; Girard-Tremblay et al., 2014; Linnman et al., 2012; Wiesenfeld-Hallin, 2005; Yu et al., 2021). Although our lab and others have observed conditioned darting in multiple experimental scenarios (Borkar et al., 2020; Colom-Lapetina et al., 2019; Greiner et al., 2019; Manzano Nieves et al., 2023; Mitchell et al., 2022; Pellman et al., 2017), the underlying biological contributors to darting are unknown. A better understanding of how brain regions known to be involved in processing aversive stimuli might differ in their recruitment during fear conditioning could give insight into the networks that drive sex differences in conditioned fear behavior.

The goals of the current study were three-fold: 1) determine if Darters exhibit higher physiological stress responses during fear conditioning; 2) determine how behavior during fear conditioning relates to behavior on other measures of aversive responding; and 3) investigate neural correlates of darting. To address these aims we first investigated rats’ change in corticosterone levels before and after fear conditioning. In another cohort of animals, we performed an acoustic startle test prior to classic fear conditioning and retrospectively compared startle responses in Darters, Non-darters, and males. In a third cohort, animals were exposed to a hot plate test one-week before fear conditioning to assess thermal pain responding in Darters, Non-Darters, and males. Finally, we quantified cFos expression following fear conditioning in brain regions known to be involved in the processing of pain and fear, including the dorsal horn of the lumbar spine (DHL) (Choi et al., 2020; Coghill, 2020; Coghill et al., 1991), the lateral parabrachial nucleus (lPbN) (Chiang et al., 2019), the sub columns of the periaqueductal gray (PAG) (Benarroch, 2012; Keay & Bandler, 2015), the lateral amygdala (LA) (Ciocchi et al., 2010; Janak & Tye, 2015; Kuner & Kuner, 2021), the central amygdala (CeA) (Ciocchi et al., 2010; LeDoux, 2000; Maren & Quirk, 2004), and the basolateral amygdala (BLA) (Corder et al., 2019; Gale et al., 2004; Gore et al., 2015). We compared activation of these regions across sexes and between Darters, Non-darters and males.

## 2. Methods

### 2.1 Subjects

All experiments were conducted in young adult (8-10 weeks) male (325-350g) and female (225-250g) Sprague Dawley rats (Charles River). Given that about 40% of females dart in a standard Pavlovian fear conditioning experiment, more females than males were used in each of the described experiments to ensure a substantial proportion of Darters to perform appropriately powered statistical analyses. Animals were same-sex, pair-housed at Northeastern University in a 12:12 light:dark cycle and had access to food and water ad libitum. Animals acclimated to the facility, undisturbed, for one week prior to testing. Testing was conducted during the light phase between the hours of 10AM and 3PM. To account for any stress that transport to and from the behavior rooms may cause, animals were placed on a rolling cart and rolled into the behavior rooms the day immediately prior to testing in addition to being handled for 5 minutes a day the two days prior to testing. All procedures were conducted in accordance with the National Institutes of Health Guide for the Care and Use of Laboratory Animals and were approved by the Northeastern University Institutional Animals Care and Use Committee.

### 2.2 Blood Collection for Corticosterone Analysis

Blood was collected immediately before and 15 minutes after fear conditioning from the lateral tail vein. Rats were placed in a restraint tube and a 25G needle was used to create a small stick hole to allow for blood effusion from the penetration point. A total volume of ∼100 μL blood sample was collected using an EDTA coated microvette tube (Kent Scientific, KMIC-EDTA). After collection, blood samples were then centrifuged at 2000 g for 20 min at 4°C. The supernatant was collected and immediately stored at −80°C until analyzed. Plasma corticosterone levels were measured using a commercially available ELISA kit (R&D Systems, KGE009) according to the manufacturer’s instructions.

### 2.3 Acoustic Startle Assay

Acoustic startle was run identical to that in Granata et al., 2022. Briefly, Med Associates’ acoustic startle hardware and software package (product number: MED-ASR-PROQ) was used in sound-proof cabinets. In each cabinet, there was an animal holder with a grid rod on top of a startle platform containing the load cell. The load cell and amplifier converted force on to the platform to the voltage representing startle response. One inch behind the animal holder, speakers delivered white noise background sound (70 decibel constant) and startle noise bursts (50 ms each at decibels of 95db, 100db, 105db, 110db, presented in random order). Sexes were counterbalanced between chambers. There were 100 trials presented in 30-second intervals. After each round, the chambers were cleaned with 70% ethanol. The experimenters involved in this assay represent people identifying as female.

### 2.4 Static Hot Plate Assay

The hot plate assay was conducted using a Corning PC 520 hot plate, heated to 52C. The temperature of the hot plate was measured before each animal underwent the assay, using an Etekcity Lasergrip 1080 infrared thermometer. The hot plate was turned on to its lowest setting and allowed to heat up for 45 minutes before a glass cylinder (12” high, 7” in diameter) was placed on top. Animals were brought into the testing room and allowed to habituate to the room for 30 minutes, after which the test began. The first rat was placed at the bottom of the cylinder and left there for 45 seconds. The built-in webcam of a 27-inch, iMac Desktop computer was used for recording the duration of the trial, recording any nociceptive behaviors the rat may display. After the rat was removed from the cylinder, the recording was stopped, and the cylinder was cleaned with 70% ethanol. The temperature of the cylinder was maintained between each rat, and if the cylinder went above desired temperature, it was removed from the plate and allowed to cool before the next rat’s test began. At the end of all the trials, the animals were placed back into the vivarium. The experimenters involved in this assay represent people identifying as either male or female.

### 2.5 Fear Conditioning

Fear Conditioning (FC) was conducted in one of four identical 20-cm^2^ chambers constructed of aluminum and Plexiglass walls (rat test cage; Coulbourn Instruments) with metal stainless steel rod flooring attached to a shock generator (Coulbourn Instruments model H13-15). The chambers were lit with a single house light, and each chamber was enclosed within a sound isolation cubicle (Coulbourn Instruments model H10-24A). An overhead, infrared digital camera recorded each trial at a frame rate of 30 frames / second. Before the animals were placed in the room for habituation, a decibel meter measured the decibel level of the CS and a shock meter measured the unconditioned stimulus (US) mA, to ensure consistency across trials. After the boxes were set up for a trial, animals were placed in the room in their cages to habituate for 30 minutes. At the start of each FC session, animals were given 5 minutes to explore the arena before the first CS-US presentation. The CS was a 30-second, 4kHz, 80dB SPL sine wave tone which co-terminated with a 0.5 second foot shock US. In experiments 1 and 2, animals were exposed to a 0.5 mA shock or no shock for a tone-only control condition (CS-only). In experiment 3, animals were assigned to a 1 mA (high) shock group, 0.3 mA (low) shock group, or a CS-only control group. There were 7 CS-US pairings throughout the trial with a mean intertrial interval of 3 minutes and a range of 1.5-5 minutes. Total test duration across all experiments was 30 minutes. After each trial, chamber walls, ceilings and the grid floors were cleaned with 70% ethanol and trays were cleaned with water and soap. Chambers were used for both male and female animals, but each test session was restricted to a single sex. Four animals ran per session, except for Experiment 3 where only two animals ran per session. The experimenters involved in this assay represent people identifying as either male or female.

### 2.6 Behavior tracking and behavioral data processing

Recorded videos from the hot plate assay were hand scored for nociceptive behaviors, such as time to lick hind paw, jumping, and fore paw attending and lifting (Carter, 1991; Minett et al., 2011). For behavior during fear conditioning, we used Noldus Ethovision software to track the animals and generate raw velocity data sheets from all video files at a sample rate of 30 frames per second. ScaredyRat, a custom python tool developed by our lab to analyze raw Ethovision data files (Mitchell et al., 2022), extracted freezing, darting, and velocity data from each animal during the tone and shock epochs. As in previous research from our lab (Gruene et al., 2015; Mitchell et al., 2022), Darters were defined as any animal that engaged in a movement across the fear conditioning chamber at or exceeding 20 cm/s in velocity during CS 3-7.

### 2.7 Euthanasia & Tissue preparation

Animals in experiments 1 and 2 were euthanized via thoracotomy. Animals in experiment 3 were anesthetized ninety minutes after the completion of fear conditioning and euthanized via transcardial perfusion with 1% saline followed by 4% paraformaldehyde (PFA) in 0.1M phosphate buffer (PBS, PH 7.4). Brains and spinal cords were extracted and post-fixed in PFA for 24 hours, and then placed in 0.1% sodium azide in PBS at 4C until slicing for storage.

### 2.8 Immunohistochemistry

All IHC procedures were conducted by an experimenter blind to the shock intensity group and Darter identity of the rats.

#### 2.8.1 Spinal cord, Amygdala, and lPbN sections: Diaminobenzadine (DAB) staining

Once ready for slicing, spinal cords and brain tissue were placed into a 30% sucrose in PBS and left in 4C until the tissue sunk to the bottom of the sucrose solution. Brain tissue was blocked prior to submersion in sucrose, while spinal cords were sectioned after. Once the tissue sank, it was embedded in Optimal Cutting Temperature (OCT) compound and frozen for slicing on a cryostat (Leica 6800) in 30 micrometer thick sections. Sections containing the central, basolateral, and lateral amygdala, the lateral parabrachial nucleus, and spinal Lumbar sections 2-4 were collected and washed in PBS three times, ten minutes each time. After the washes, endogenous peroxidase activity was quenched for 15 minutes in a 0.5% hydrogen peroxide solution in PBS, after which the slices were mounted into microscope slides. Once sufficiently dry, slides were rinsed three times with PBS, ten minutes each time. All washes were performed on a shaker and all incubations took place in an opaque incubation chamber, both at room temperature. They were then blocked in a 10% Normal Goat Serum (NGS, S26, Millipore) in 0.1% PBS-T for one hour at room temperature. Following blocking, slides were incubated in a Rabbit anti cFos primary antibody (Rabbit IgG Fos Abcam – ab190289) with 10% NGS in 0.1% PBS-T, overnight at room temperature. Eighteen to twenty-four hours later, slides were rinsed in PBS for ten minutes, three times before secondary antibody incubation in biotinylated goat anti-rabbit (PK-6101, 1:200) in 1.5% NGS in PBS for 2 hours. An ABC solution was prepared using the Vectastain Elite ABC Kit (Vector Laboratories) 30 minutes prior to use to allow complex to form. After secondary antibody incubation, slides were rinsed three times, five minutes each in PBS and then incubated for 45 minutes in the ABC solution. After ABC, slides were rinsed three times, five minutes each, in 0.1M phosphate buffer and then incubated in DAB solution. The spinal cord and amygdala sections incubated for 20 minutes and the lPbN incubated for 2 minutes, due to differences in staining intensity. After incubation, slides were rinsed in PBS three times, five minutes each and then dried and cover-slipped for imaging. Images were taken at 10X magnification and stitched together using a Keyence BZX 710 at 1/500s exposure.

#### 2.8.2 Fluorescent Immunohistochemistry

Fifty micrometer PAG sections were collected using a vibrating microtome (Leica VT 1000S). PAG sections –8.04 from Bregma from each animal were isolated and washed three times for ten minutes each in 0.1% Triton-X 100 in PBS (PBS-T), and then incubated in a blocking buffer containing 10% Normal Donkey Serum (NDS) in PBS-T for 1 hour at room temperature. Sections were then incubated overnight at 4C in a polyclonal rabbit anti-cfos 1:2000 (ABCam AB190289). Eighteen to twenty-four hours later, slices were rinsed in 0.1% PBS-T three times, ten minutes each, before incubating for 1.5 hours in a donkey anti-mouse secondary antibody 1:1000 Alexa 647 (Jackson Immuno product #715-605-150) at room temperature. After secondary incubation, sections were rinsed three times, ten minutes each in 0.1% PBS. At the conclusion of the washes, sections were mounted on microscope slides, cover slipped, and dried overnight. Slides were imaged at 10X magnification on the same Keyence as DAB-stained sections and the PAG sections were imaged in a 3×3 grid at 3.5s exposure.

### 2.9 Cell quantification

Images from the brain and spinal cord were imported to ImageJ software and number of cFos+ cells were quantified using a customized cell counting macro by an experimenter blind to animal shock intensity condition and Darter identity. We used one section per region, as regional functions can vary on a rostral / caudal gradient. We chose to focus on sections that were previously identified as being involved in pain or fear processing. Sections used were as follows: laminae i-iii in lumbar spinal cord sections 2-4 (Westlund & Willis, 2015), lateral parabrachial nucleus bregma –8.76 (Raver et al., 2020), periaqueductal gray columns bregma –8.04 (Carrive et al., 1997), central amygdala bregma –2.40 (Prusator & Greenwood-Van Meerveld, 2017), lateral amygdala bregma –2.76 (Swanson & Petrovich, 1998), basolateral amygdala bregma –3.0 (Vazdarjanova & Mcgaugh, 1999).

### 2.10 Statistical Analysis

All statistical analyses were conducted using GraphPad Prism software version 10.2.3 (one and two-way ANOVAs, chi-squared tests) and SPSS version 28 (three-way ANOVAs and correlation matrices). When appropriate, post hoc tests (Sidak’s multiple comparisons, Dunnett’s multiple comparisons, or Tukey’s) were used on main effects and interactions. In standard cohorts, there are not sufficient N’s to run statistics on male Darters so they are not split by Darter identity. Outliers were removed if they were +/− two standard deviations away from a group’s mean and were calculated for each individual group analyzed (e.g. 0.3 mA females, 0.3 mA female Non-darters, 0.3 mA female Darters: outliers were calculated at each level).

In experiment 1, a 2×2×2 mixed model three-way analysis of variance (ANOVA) was performed to examine between subject factors of testing condition (shock exposure or CS only) and sex (male, female) and the within subject factor of time (CORT levels pre and post fear conditioning). For females, a 2×2 mixed model ANOVA was performed on the between subject factor of Darter identity (Darter, Non-darter) and the within subject factor of time (CORT levels pre and post fear conditioning).

In experiment 2, a 3×4 mixed model, two-way ANOVA was conducted with the between subject factor of Darter identity (Darter, Non-darter, males) and decibel intensity (95, 100, 105, 110 dB) for both peak startle amplitude and latency to startle.

In experiment 3, for nociceptive behavior, a one-way ANOVA was conducted on the between subject factor of Darter identity (Darter, Non-darter, male) for hind withdraw and fore paw attending. A 3×2 mixed model, two-way ANOVA was conducted on the between subject factor of Darter identity (Darter, Non-darter, male) and the within subject factor of paw withdraw latency (hind paw, fore paw). For fear conditioning behavior, chi-squared tests were conducted to determine the differences in Darter percentages between shock intensity groups in males and females. A 2×3×7 mixed-model, three-way ANOVA was conducted on the between subject factors of sex (male, female) and shock intensity (CS only, 0.3 mA, 1 mA) and the within subject factors of tone or shock (tones / shocks 1 thru 7), on percent of time spent freezing and shock response (cm/s). In females, a 3×2×7, mixed model 3-way ANOVA of the between subject factors of shock intensity (CS only, 0.3 mA, 1 mA), Darter identity (Darter, Non-darter), and tone or shock (tones / shocks 1 thru 7), was conducted for freezing and shock response.

To avoid cohort effects for cFos cell counts in experiment 3, we normalized data to each cohort’s control condition (CS only) by taking the average of the controls from each cohort, and then dividing each individual animals’ counts by their respective cohort’s control average. For each region, 2×2 between-subjects ANOVAs (shock intensity, 0.3 mA and 1 mA and sex, male and female) were conducted on cFos expression in regions of interest. Females were further broken down and 2×2 between-subjects ANOVAs (shock intensity 0.3 mA and 1 mA and Darter identity, Darter, Non-Darter), were conducted on cFos expression in the same regions. Pearson’s bivariate correlations of activity between the regions of interest were conducted.

## 3. Results

### 3.1 Experiment 1

A graphical representation of experiment 1 can be found in Figure 1A. Twenty-seven females (19 exposed to CS plus shock, 8 CS-only controls) and 17 males (10 shock, 7 controls) were used in this experiment.

**Figure 1.**
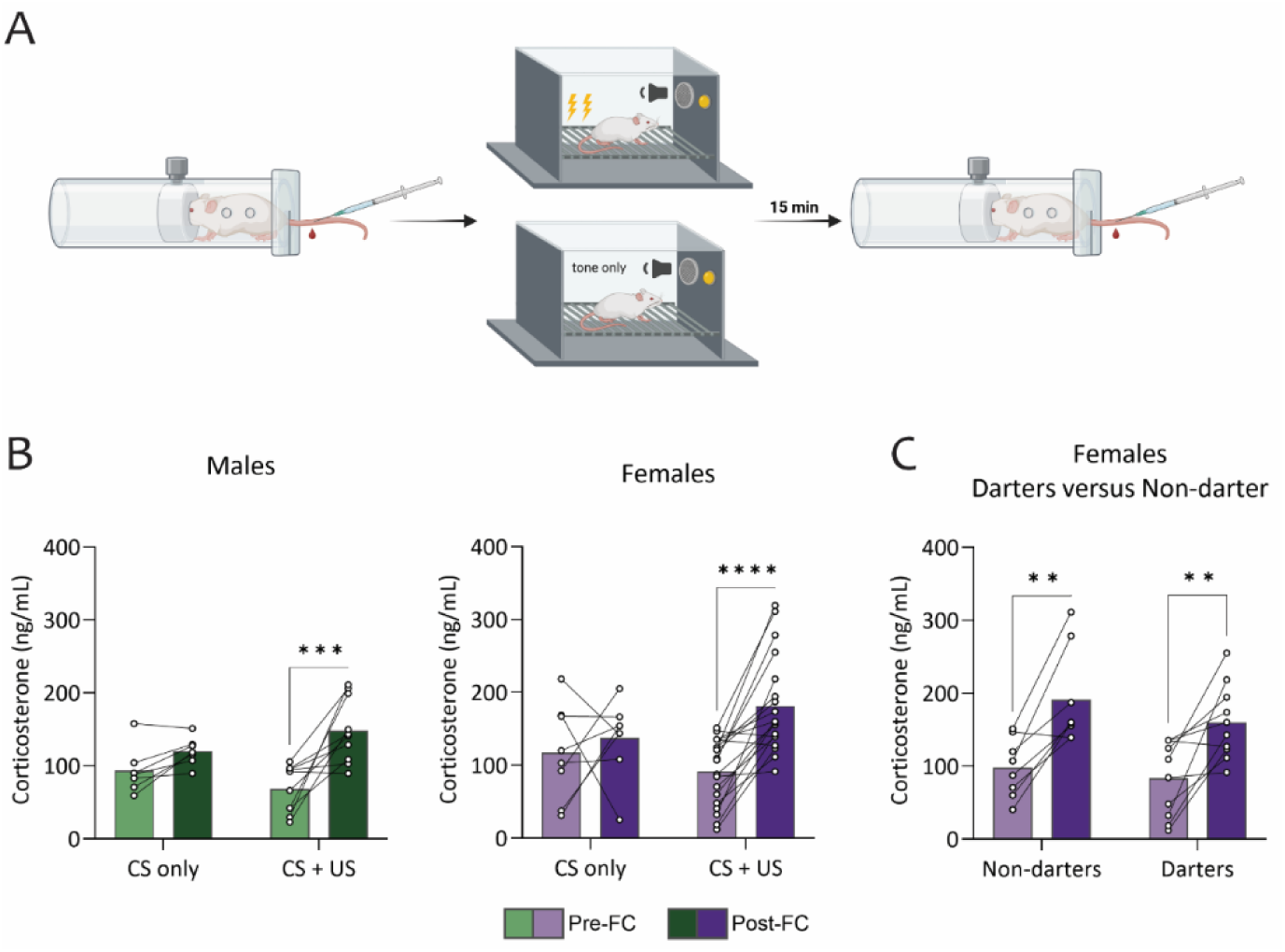
Fear conditioning induces comparable increases in corticosterone levels in all animals. **A**. Experimental design consisted of blood plasma samples taken immediately before and 15 minutes after a 7 CS-US fear conditioning paradigm. **B**. Corticosterone levels increased in fear conditioned males and females, but not in CS-only males and females. **C**. Corticosterone levels increased in all Darters and Non-darters following fear conditioning. (**) p < 0.01, (***) p < 0.001, (****) p < 0.00001. Design figure in panel A created in Birorender.

#### 3.1.1 Corticosterone levels do not differ between Darters and Non-darters

Measurements of blood plasma levels of corticosterone taken immediately before and 15 minutes after fear conditioning show that corticosterone levels increased in response to fear conditioning in males and females (Fig. 1B). CS-only controls did not show an increase in corticosterone levels (Fig 1B). A mixed model ANOVA comparing the between subjects factors of shock exposure and sex with the within subject factor of pre and post conditioning corticosterone levels (pre/post timing) found a significant main effect for pre/post timing (F(1, 31) = 13.95, p = 0.0008). There was also a significant interaction between pre/post timing and shock exposure (F(1,31) = 5.72, p = 0.023), Sidak’s multiple comparison test showed a significant increase in CORT levels following fear conditioning in the shock exposure group in both males (p = 0.01) and females (p = 0.006). No difference was found in the groups exposed to CS-only. Because corticosterone only increases in the fear conditioned group, this validates that increases in corticosterone levels following fear conditioning are indeed due to shock exposure and not stress from manipulation/testing.

Of the fear conditioned females, eight were Darters and ten were Non-darters. In both female Darters and Non-darters, corticosterone levels increased from baseline following fear conditioning (Fig. 1C). A mixed model ANOVA comparing the between subject factor of Darter identity (Darter, Non-darter) and the within subject factor of pre and post conditioning corticosterone levels in females found a significant main effect for pre/post timing (F(1,16) = 34.09, p < 0.0001). Sidak’s multiple comparisons showed significant differences for Darters (p = 0.0023) and Non-darters (p = 0.0011). There was no main effect of Darter identity.

### 3.2 Experiment 2

A graphical representation of experiment 2 can be found in Figure 2A. Twenty-four females and 19 males were used in this experiment.

**Figure 2.**
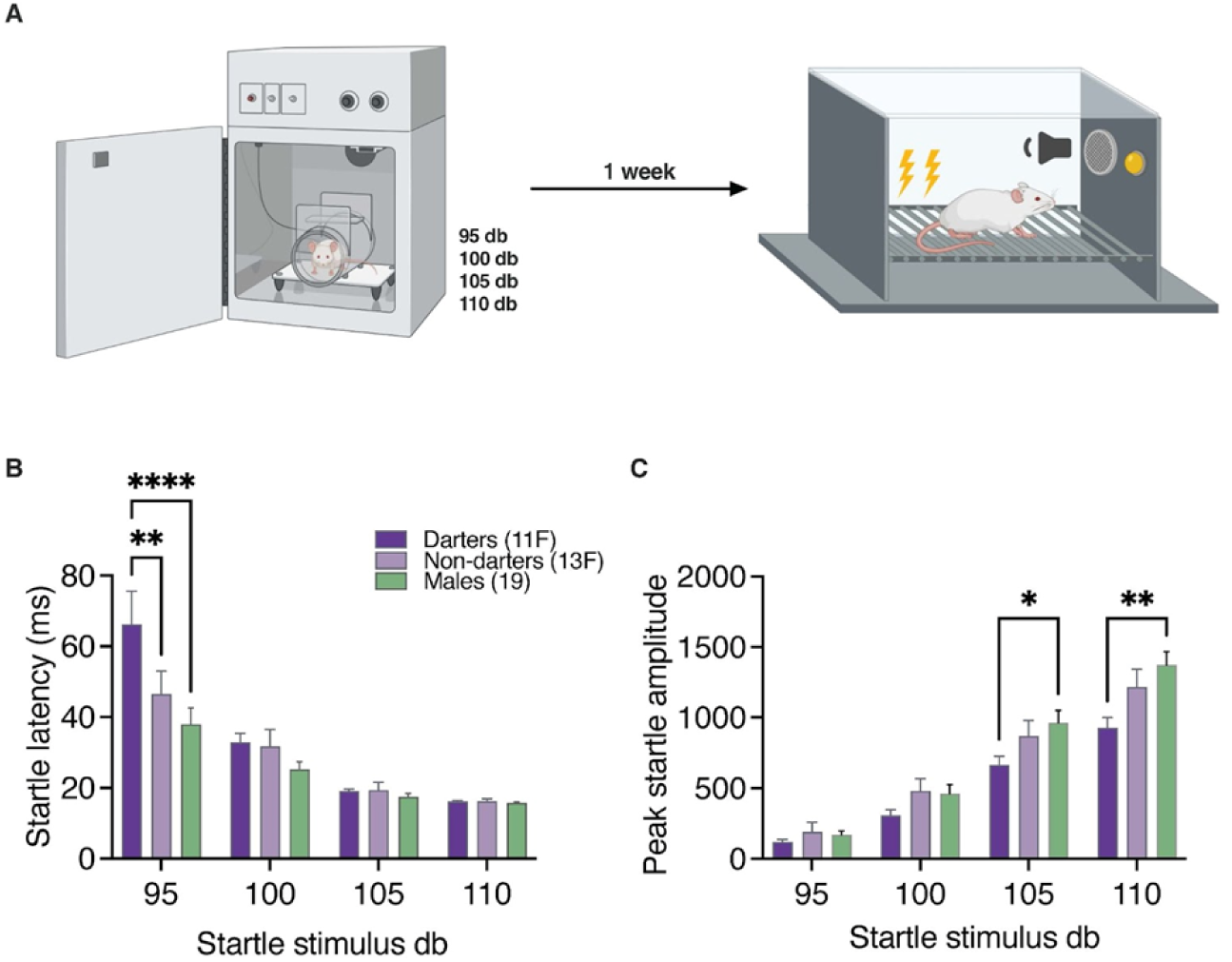
Darters do not exhibit enhanced startle responses. **A**. Experimental design consisted of the acoustic startle test where animals were exposed to a series of startle-evoking white noises at four different decibel levels presented in a random order for one hour. One week later, animals ran through a 7 CS-US fear conditioning paradigm. **B**. Darter females have a greater latency to startle than non-darter females and males at 95 db stimuli. **C**. Males have a greater peak startle amplitude than darter females at 105 db and 110 db stimuli. (*) p < 0.05, (**) p < 0.01, (****) p < 0.00001. Design figure in panel A created in Birorender.

#### 3.2.1 Darters do not have greater startle responses than Non-darters and males

A mixed model ANOVA comparing the between subject factor of Darter identity and the within subject factor of stimulus decibel on startle latency found significant main effects for both Darter identity (F(2, 160) = 6.582, p = 0.0018) and decibel (F(3, 160) = 51.48, p < 0.0001) (Fig 2B). There was also a significant interaction between Darter identity and decibel (F(6, 160) = 3.164, p = 0.0058). Dunnett’s multiple comparisons test found significant differences between female Darters and Non-darters (p = 0.0013) and female Darters and males (p < 0.0001) only at the 95 decibel stimulus. In the acoustic startle test, at lower (95dB) decibels, female Darters had a longer latency to startle than female Non-darters and males, but at higher decibels (100 dB, 105 dB, 110 dB), female Darters did not differ from female Non-darters and males. Together, these data show that at lower decibels, Darters are slower to startle than Non-darters and males.

A two-way ANOVA comparing the between subject factor of Darter identity and the within subject factor of stimulus decibel on peak startle amplitude found significant main effects for decibel (F (1.737, 69.46) = 290.1, p < 0.0001), a trending main effect for Darter identity (F(2, 40) = 2.748, p = 0.07), and a significant interaction between Darter identity and decibel (F(6, 120) = 3.822, p = 0.0016) (Fig 2C). Dunnett’s multiple comparisons test found significant differences between males and female Darters at 105 decibels (p = 0.027) and 110 decibels (p = 0.002). The lack of a difference between Darters’ and Non-darters’ peak startle suggests that Darters are not merely hyperreactive and more sensitive to a broad range of aversive stimuli compared to Non-darters.

### 3.3 Experiment 3

A graphical representation for experiment 3 can be found in Figure 3A. Sixty-two females (26 0.3 mA, 20 1 mA and 16 CS-only) and 59 males (22 0.3 mA, 21 1 mA, and 16 CS-only) went through the hot plate assay followed by fear conditioning approximately one week later and then cFos analysis. Twenty-four males and 22 females were included in the final hot plate analysis and the rest were excluded due to a video quality issue that prevented accurate scoring. For the cFos portion of the experiment, each region has different N’s for males and females due to timing effects of staining.

**Figure 3.**
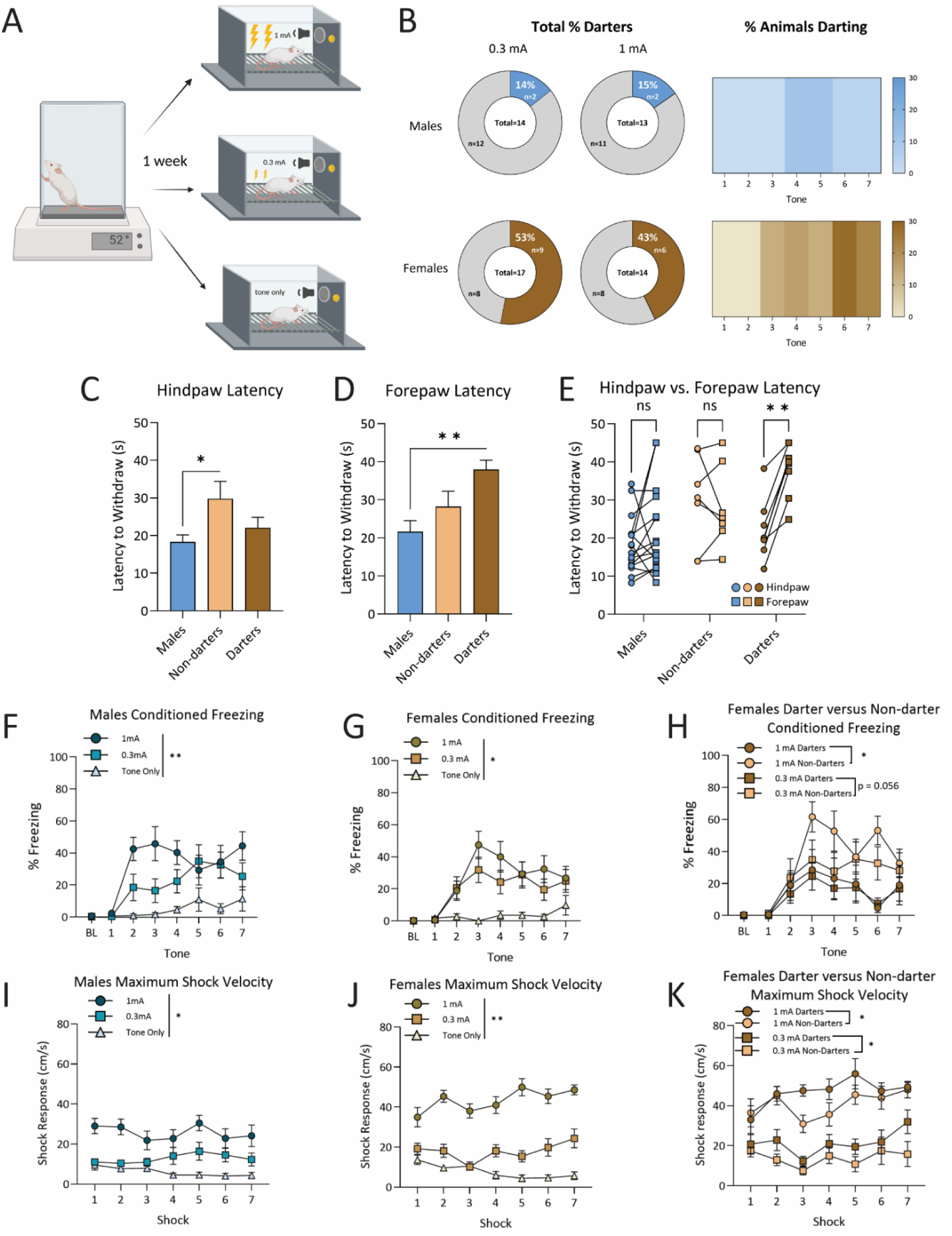
Behavior results for Experiment 3. **A**. Experimental design consisted of one day of the hot plate assay to obtain baseline thermal pain threshold. One week following exposure to the hot plate, animals ran through a 7 CS-US fear conditioning paradigm in one of three groups: 1mA shock, 0.3mA shock, or CS-only control groups. **B.** Proportion of Darters in males and females across 0.3 and 1 mA shock intensity groups and heatmaps for both sexes showing prevalence of conditioned darting across fear conditioning tone **C.** Female Non-darters have a longer latency to withdraw their hind paw than males. **D.** Female Darters have a longer latency to withdraw their forepaw than males. **E.** Darters consistently withdraw their hind paws before their forepaws, a pattern not seen in Males or non-Darters. **F.** Percent of time spent freezing increased because of increasing shock intensity in males **G.** Percent of time spent freezing increased with higher shock intensity in females. **H.** In females, non-Darters froze more than Darters, regardless of shock intensity. **I.** Shock response increased with higher shock intensity in males **J.** Shock response increased with higher shock intensity in females. **K.** In Females, Darters showed higher shock response than non-Darters exposed to the same shock intensity. (*) p < 0.05, (**) p < 0.01. Design figure in panel A created in Birorender.

#### 3.3.1 Behavior comparisons across hot plate and fear conditioning paradigms

#### 3.3.2 There is a relationship between nociceptive behavior and Darter identity

During fear conditioning, females darted more than males, as shown by a significant chi-squared test (χ^2^(1)=7.384, p = 0.003) and darted to the later tones (Fig 3B). Non-darter females had a longer latency to withdraw their hind paws than males (Fig 3C). One-way ANOVA revealed a significant effect of Darter identity on hind paw withdrawal (F(2, 28) = 4.17, p = 0.03), and Dunnett’s multiple comparison tests resulted in a significant difference between males and Non-darters (p = 0.02), but not males and Darters (p = 0.59). A one-way ANOVA similarly revealed a significant effect of sex on forepaw attending latency (F(2,28) = 6.60, p = 0.005) (Fig 3D).

Dunnett’s multiple comparisons revealed that males still showed the quickest latency to attend, but only in comparison to female Darters (p = 0.003), who had the longest latency to attend, and that males were not different than not Non-darters (p = 0.36). Finally, we examined within-subjects latencies between hind and fore paw. A two-way ANOVA revealed a main effect of Darter identity (F(2,56) = 7.699, p = 0.001), paw (hind vs. fore) (F(1,56) = 5.048, p = 0.029) and a significant interaction between Darter identity and paw (F(2,56) = 3.47, p = 0.038). Darters were the only group to show a significant within group difference between fore and hind paw latency, consistently withdrawing their hind paw before their fore paw (p = 0.002), as identified by a post-hoc Tukey’s test, corrected for multiple comparisons (Fig 3E). Therefore, unlike males and Non-darters, Darters exhibit a predictable pattern of paw withdrawal order in a hot plate test.

#### 3.3.3 Conditioned and unconditioned responses during fear conditioning differ by shock intensity and sex

Both males and females froze more to higher shock intensities, and female Darters froze less than their Non-darter counterparts. A 3-way ANOVA of sex, shock intensity, and tone revealed a significant main effect of shock intensity on freezing (F(2, 79) = 18.834, p < 0.001), in that males and females in either shock group froze more than CS-only animals (p < 0.05 for both sexes when 0.3 and 1 mA are compared to CS-only) (Fig 3F,G). When females were split by Darter identity, a 3-way ANOVA of shock intensity, Darter identity and tone revealed a significant main effect of Darter identity on freezing (F(1,31) = 10.15, p = 0.003): 1 mA Darters froze less than their Non-darter counterparts (p = 0.018), and 0.3 mA Darters trended towards freezing less than their Non-darter counterparts (p = 0.056) (Fig 3H).

Both males and females reached higher maximum shock velocities in response to higher shock intensities, and female Darters in both shock intensities reached higher shock velocity than their Non-darter counterparts. A 3-way ANOVA revealed that animals exposed to higher shock intensities reached higher maximum shock velocities (F(2,79) = 66.37, p < 0.001), regardless of sex (females and males, all p < 0.05) (Fig 3I,J). There was also a main effect of sex (F(1,79) = 14.91, p < 0.001), and females had overall higher shock response than males at the 1mA shock intensity (p < 0.001). When split by Darter identity, a 3-way ANOVA revealed that in females there was a significant effect of shock intensity (F(1, 31) = 84.23, p < 0.001) and Darter identity (F(1,31) = 9.28, p = 0.005) on shock response. Female Darters at both 0.3 and 1 mA shock intensities reached higher maximum shock velocities than their Non-darter counterparts (p < 0.05 for both shock intensities) (Fig 3K).

#### 3.3.4 cFos expression in key regions following fear conditioning

The experimental design for all c-fos assays is shown in Figure 4A. Animals were excluded from specific regions if tissue from that region was too damaged to accurately count.

**Figure 4.**
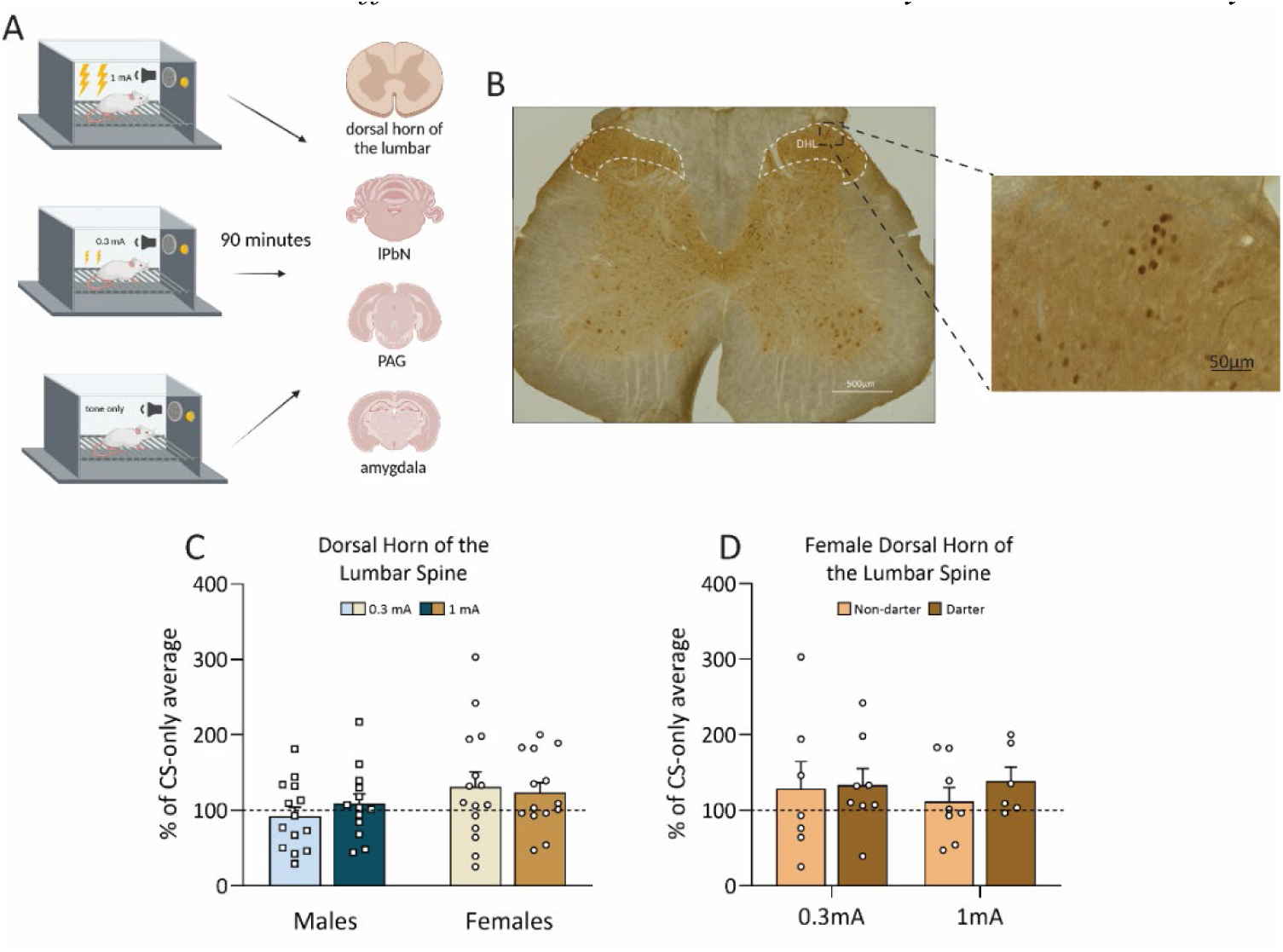
cFos activity in the dorsal horn of the lumbar spine following fear conditioning. **A**. Experimental design for cFos portion of the experiment. **B.** Representative image of L3 of the Dorsal Horn of the Lumbar stained with DAB for cFos. **C.** There were no differences in cFos expression across shock intensities or sexes. **D.** There were no differences in cFos expression across conditioned responses. Dotted line indicates CS-only animal cFos levels. Design figure in panel A created in Birorender.

#### 3.3.5 cFos expression in the DHL did not differ based on sex, shock intensity, or Darter identity

Thirty-nine female (16 0.3 mA, 14 1 mA, 9 CS-only) and 37 male (13 0.3 mA, 14, 1 mA, 10 CS-only) rats were included in the DHL analyses. Figure 4B shows a representative image of cFos DAB staining in the DHL. A two-way ANOVA revealed that there were no significant effects of shock intensity (F(1,52) = 0.085, p = 0.77), or sex on cFos expression, although females trended towards more cFos expression, regardless of shock intensity (F(1,52) = 3.161, p = 0.08) (Fig 4C). When females were separated by Darter identity, a two-way ANOVA revealed no effect of Darter identity (F(1,25) = 0.379, p = 0.53) or shock (F(1,25) = 0.174, p = 0.82) (Fig 4D), suggesting that the potential increase in cFos expression seen in females is not due to any differences between Darters and Non-darters in the DHL.

#### 3.3.6 cFos expression in the lPbN is sex and shock intensity-dependent

Twenty-two female (7 0.3 mA, 8 1 mA, 7 CS-only) and 23 male (7 0.3 mA, 8 1 mA, 8 CS-only) rats were included in the lPbN analyses. Figure 5A shows a representative image of cFos DAB staining in the lPbN. A two-way ANOVA revealed a significant main effect of shock intensity (F(1,19) = 6.199, p = 0.022), and Sidak’s multiple comparisons post hoc test revealed that this was driven by females’ heightened expression in the 1mA group compared to the 0.3 mA group (p = 0.011) (Fig 5B). There was also a significant main effect of sex on cFos expression (F(1,19) = 7.843, p = 0.011) and Sidak’s multiple comparisons post hoc test revealed that this was driven by the 1 mA group, where females showed significantly more expression than males (p = 0.011) (Fig. 6B). When females were grouped by Darter identity, a two-way ANOVA showed that the effect of shock intensity almost reached significance (F(1,10) = 4.653, p = 0.056), but there was no effect of Darter identity (F(1,10) = 0.082, p = 0.78) (Fig 5C). The increase in cFos expression in the 1 mA females compared to the 0.3 mA females was not due to Darter identity.

**Figure 5.**
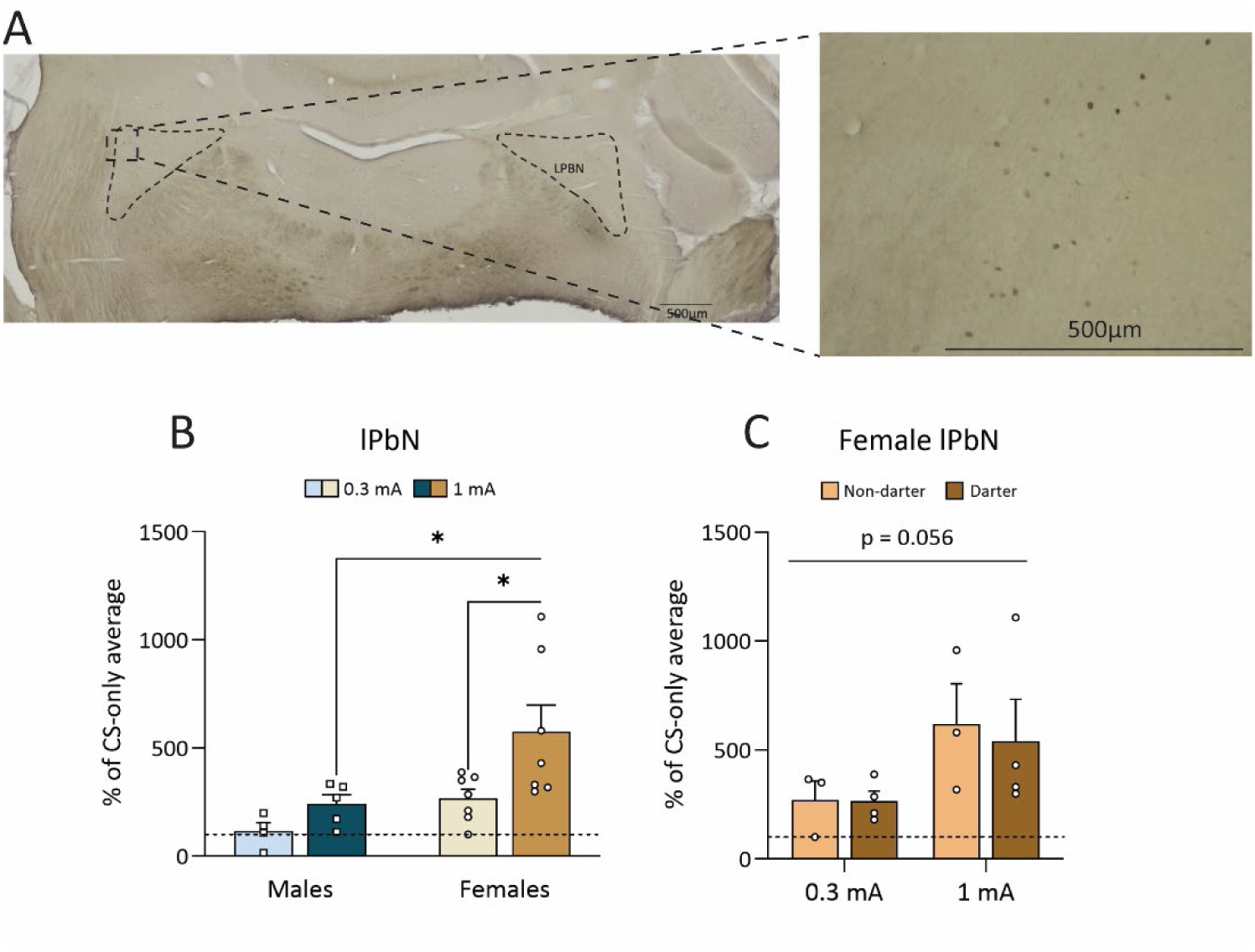
cFos activity in the lateral parabrachial nucleus following fear conditioning. **A**. representative image of the lPbN following staining. **B.** There was as significant effect of shock in females only, and a significant effect of sex, driven by the 1 mA animals. **C.** When females were separated by conditioned response, 1 mA animals trended towards higher cFos levels, regardless of CR. (*) p < 0.05. Dotted line indicates CS-only animal cFos levels.

**Figure 6.**
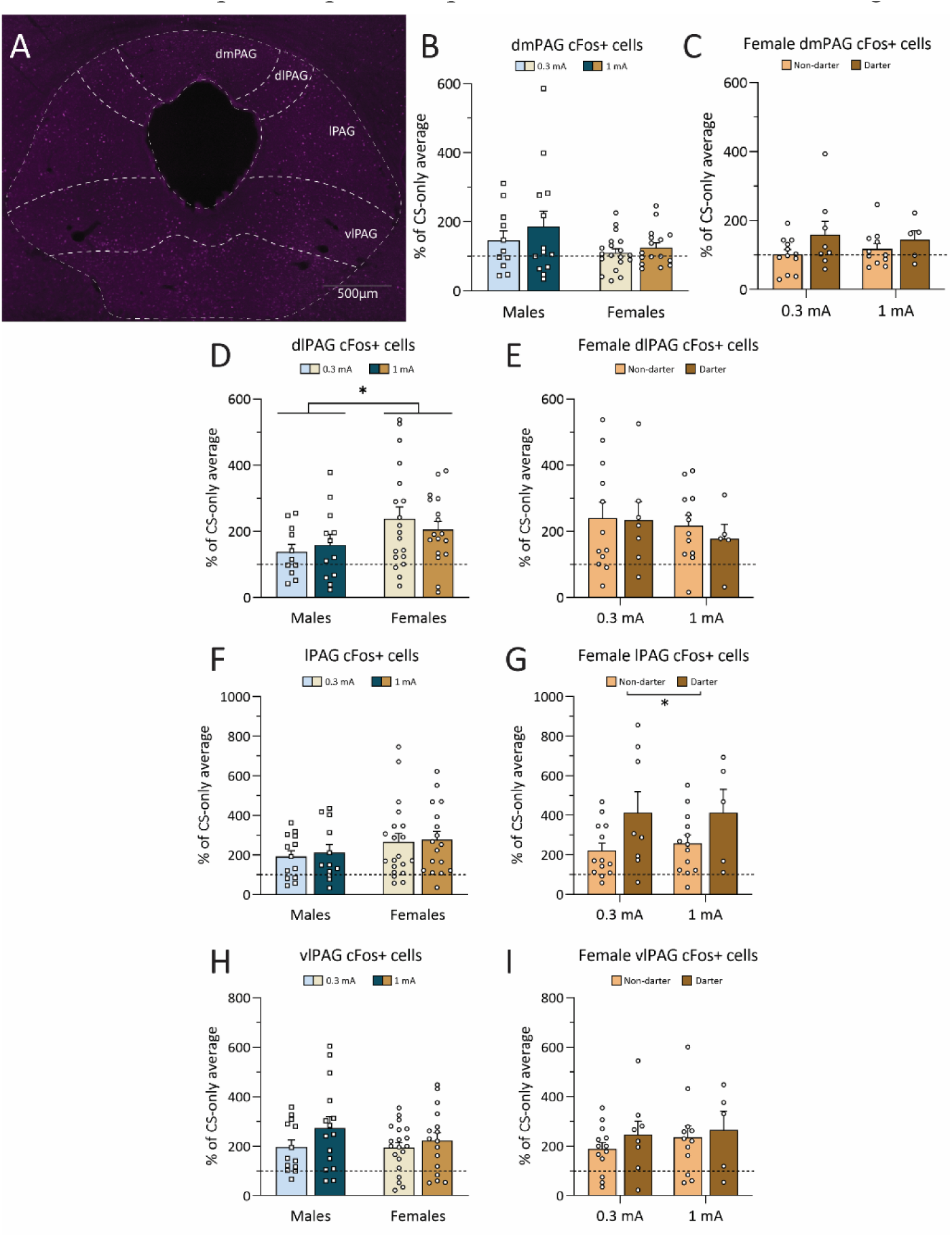
cFos analyses from the columns of the periaqueductal gray following fear conditioning. **A**. Representative image of the immunofluorescent staining for cFos in the PAG. Image has been edited for this figure to make cells more visible to the naked eye. **B-C.** cfos+ cells in the dmPAG, by sex and for females only by conditioned response. No differences were found between any groups. **D.** There was a significant effect of sex in the dlPAG: Females had more cFos expression than males. **E.** There was no significant effect of shock or conditioned response in the dlPAG. **F.** No sex or shock effect was found in the lPAG. **G.** There was a significant effect of conditioned response in the lPAG: Darters showed increased cFos+ cells in the lPAG compared to Non-darters, regardless of shock intensity. **H.** There was no effect of shock or sex in the vlPAG. **I.** There was no effect of conditioned response in the vlPAG. (*) p < 0.05. Dotted line indicates CS-only animal cFos levels.

#### 3.3.7 cFos expression in the PAG following fear conditioning varies depending on column

Figure 6A is a representative image of each column of the PAG. Fifty-one female (24 0.3 mA, 19 1 mA, 8 CS-only) and 35 male (13 0.3 mA, 14, 1 mA, 8 CS-only) rats were included in the PAG analyses. The PAG was analyzed based on sub columns, as each column has unique roles in pain and fear responding (Deng et al., 2016; Keay & Bandler, 2015; Reis et al., 2023; Vianna et al., 2001; Watson et al., 2016).

A two-way ANOVA showed no effect of shock intensity (F(1,55) = 1.193, p = 0.28) in the dorsomedial (dmPAG), but there was a trending effect of sex (F(1,55) = 3.639, p = 0.062), with males appearing to have higher cFos expression than females (Fig 6B). A two-way ANOVA showed no effect of Darter identity in females (F(1,32) = 0.0005, p = 0.98) (Fig 6C). In the dorsolateral (dlPAG), a two-way ANOVA showed no significant effect of shock intensity (F(1,55) = 0.024, p = 0.85), but there was a significant effect of sex (F(1,55) = 5.047, p = 0.029) and Sidak’s multiple comparisons post hoc test revealed a trending effect for females in the 0.3 mA group compared to males in the 0.3 mA group (p = 0.067) (Fig. 6D). When females were split by Darter identity, a two-way ANOVA showed no effect of Darter identity (F(1, 32) = 0.670, p = 0.42) (Fig 6E). These results from the dorsal columns of the PAG suggest that the dmPAG is not heavily recruited by either sex during fear conditioning and that the dlPAG is recruited in females, but not in males.

A two-way ANOVA revealed that there was no significant effect of shock intensity in the lPAG (F(1,58) = 0.144, p = 0.70), or of sex F(1,58) = 2.74, p = 0.10), (Fig 6F). When separated by Darter identity, a two-way ANOVA showed that there was a significant effect of Darter identity on cFos expression (F(1,35) = 6.236, p = 0.02). This failed to reach significance in post-hoc analyses, but 0.3 mA Darters trended to have more expression than their Non-darter counterparts, regardless of shock intensity (p = .08) (Fig 6G). Overall, this suggests that recruitment of the lPAG is higher in Darters compared to Non-darters. No differences were found across sexes or Darter identity in the vlPAG. As analyzed in a two-way ANOVA, the vlPAG did not show a significant effect of shock intensity (F(1,60) = 2.69, p = 0.11) or sex (F(1,60) = 0.711, p = 0.40) (Fig 6H). No effect was found when females were analyzed by Darter identity via two-way ANOVA (F(1,34) = 0.484, p > 0.05) (Fig 6I). It is important to note that cFos expression in the dlPAG, lPAG and vlPAG in males and females at both shock intensities was well above the CS-only mean (dotted lines on all graphs).

#### 3.3.8 cFos expression in the lateral, central and basolateral amygdala varies based on shock intensity and Darter identity

Thirty-four female (14 0.3 mA, 11 1 mA, 9 CS-only) and 37 male (13 0.3 mA, 14, 1 mA, 10 CS-only) rats were included in the amygdala analyses. Figure 7A is a representative image of DAB cFos staining in the LA. A two-way ANOVA revealed no significant effect of shock intensity (F(1,37) = 0.378, p = 0.54) or sex (F(1,37) = 0.154, p = 0.70) (Fig 7B). When females were separated by Darter identity, there was a significant shock intensity x Darter identity interaction (F(1,16) = 5.06, p = 0.04) and significant main effect of Darter identity (F(1,16) = 13.69, p = 0.002) (Fig 7C). Sidak’s multiple comparisons post-hoc test revealed that Darters had higher cFos expression than Non-darters in the 1mA condition (p = 0.002).

**Figure 7.**
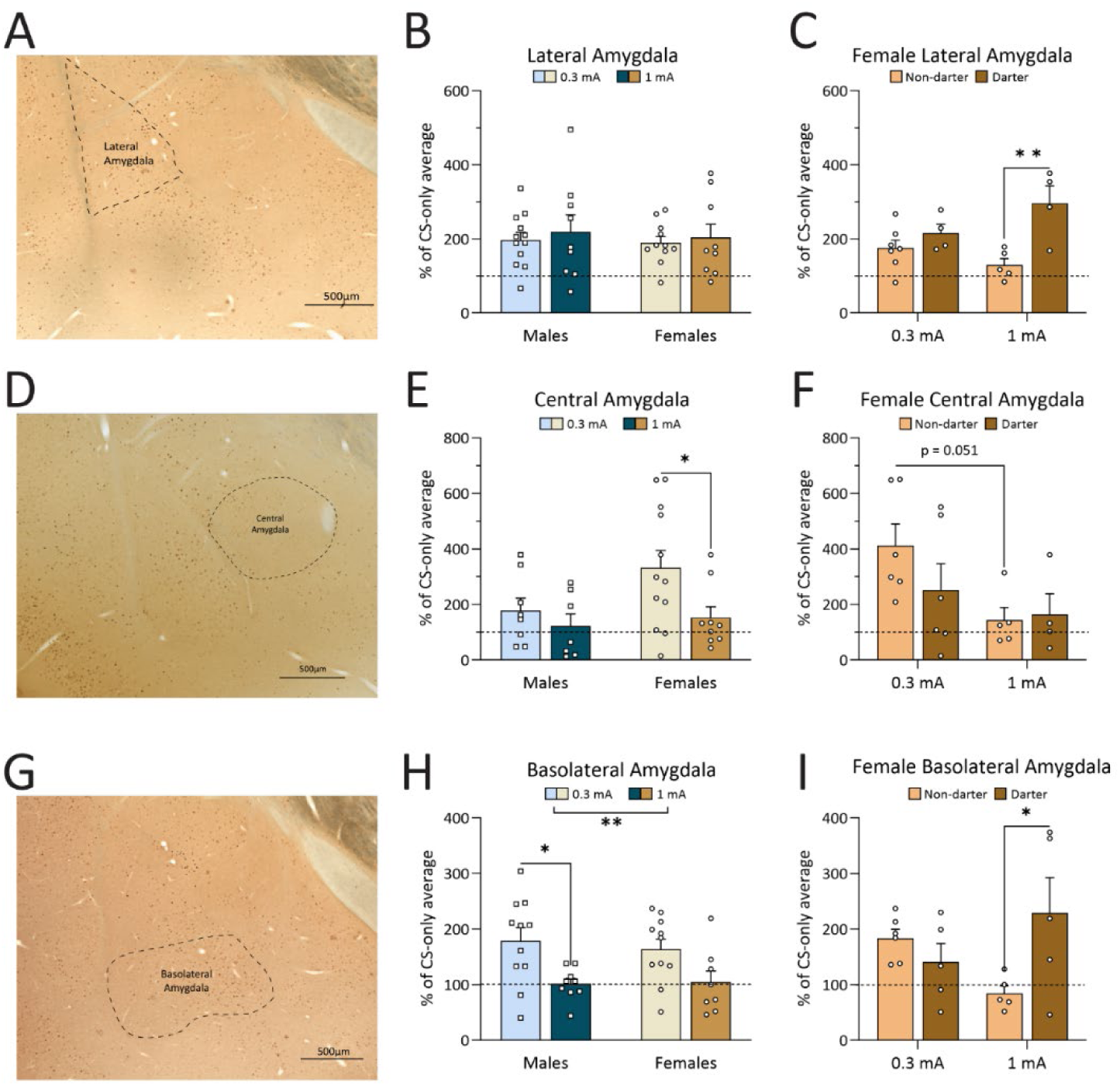
cFos analyses from the lateral, central, and basolateral amygdala. **A**. Representative image of DAB cFos staining in the lateral amygdala. –2.76 from Bregma. **B.** There was no main effect of shock intensity or sex in the lateral amygdala. **C.** There was a significant effect of shock intensity and a significant interaction between conditioned response and shock intensity in females: At higher shock intensities, female Darters had more cFos expression in the lateral amygdala than Non-darters. **D.** Representative image of DAB cFos staining in the central amygdala. –2.40 from Bregma. **E.** No effects of shock intensity or sex were found in the central amygdala. **F.** There was a significant shock x conditioned response interaction in females: Female Non-darters had significantly less cFos expression in the central amygdala at higher shock intensities; Darter did not follow this pattern. **G.** Representative image of DAB cFos staining in the basolateral amygdala. –3.00 from Bregma. **H.** There was a significant effect of shock intensity in the BLA, although post-hoc tests did not reach significance. **I.** There was a significant shock x conditioned response interaction in females: Non-darters decreased cFos expression at higher shock intensities while Darters increased. (*) p < 0.05, (**) p < 0.01, (***) p < 0.001. Dotted line indicates CS-only animal cFos levels.

Figure 7D is a representative image of DAB cFos staining in the CeA. A 2-way ANOVA revealed a significant main effect of shock intensity (F(1,32) = 4.651, p = 0.039), and Sidak’s multiple comparison tests revealed that the 0.3 mA females had significantly more expression than the 1 mA females (p = 0.033). There was no significant effect of sex on cFos expression (F(1,32) = 3.866, p = 0.10) (Fig 7E). When females were separated by Darter identity there was a significant effect of shock intensity (F(1,17) = 4.95, p = 0.0399) (Fig 7F). Sidak’s multiple comparisons test revealed that Female Non-darters were just shy of having significantly more cFos expression at 0.3 mA than 1 mA (p = 0.051), and Darters did not differ in their expression (p = 0.70).

Figure 7G is a representative image of the BLA with DAB cFos staining. A two-way ANOVA revealed no significant main effect of sex (F(1,35) = 0.09995, p = 0.75), but a significant main effect of shock intensity (F(1,35) = 12.58, p = 0.001) (Fig 7H), with both males and Non-darter females showing increased cFos expression in response to 0.3 mA shocks. Sidak’s multiple comparisons revealed that 0.3 mA males had significantly more cFos expression than 1 mA males (p = 0.014) and the same effect was trending in females (p = 0.075). When females were separated by Darter identity, a significant shock intensity x Darter identity interaction was found (F(1,17) = 6.87, p = 0.018) (Fig 7I). Sidak’s multiple comparisons post hoc test revealed that, similar to the LA, Darters had higher cFos expression than Non-darters at 1 mA (p = 0.02).

#### 3.3.9 Patterns of cFos expression varied based on sex and shock intensity

We next asked whether cFos expression across subregions reflected coordinated network activity, and whether these networks shifted as shock intensity increased. To explore these relationships, we conducted Bivariate Pearson’s correlations of cFos expression between all subregions, among all experimental groups. Supplementary tables 1-4 (Appendix A) show all the correlations between the DHL, lPbN, the sub columns of the PAG and the lateral, basolateral and central amygdala. Females were not further split by Darters and Non-darters because there were not enough animals (N < 10 for all regions except the PAG) in each group to perform correlations, once animals were split by shock intensity, Darter identity, and loss of tissue was considered.

Figure 8A shows inter-region correlations for males at each shock intensity. In 0.3 mA males, the dlPAG, lPAG and vlPAG were positively correlated with each other, and the dmPAG was only positively correlated with the vlPAG. In 1 mA males, all the sub columns of the PAG (including the dmPAG) were positively correlated with each other, and the vlPAG also showed a positive correlation with the LA. Figure 8B shows correlations among females at each shock intensity. In 0.3 mA females, all the sub columns of the PAG were significantly positively correlated with each other, and the CeA and LA were negatively correlated with each other. This pattern changed in 1 mA females. Only the vlPAG and lPAG were positively correlated with each other. The BLA was negatively correlated with the vlPAG and lPAG and the CeA was positively correlated with the lPAG. Correlations between the CeA, BLA, vlPAG and lPAG all have sample sizes of less than 10. Females exposed to a higher shock intensity lost the tight interconnectivity between the PAG sub columns, which contrasts with males exposed to a higher shock intensity, who maintain their PAG interconnectivity.

**Figure 8.**
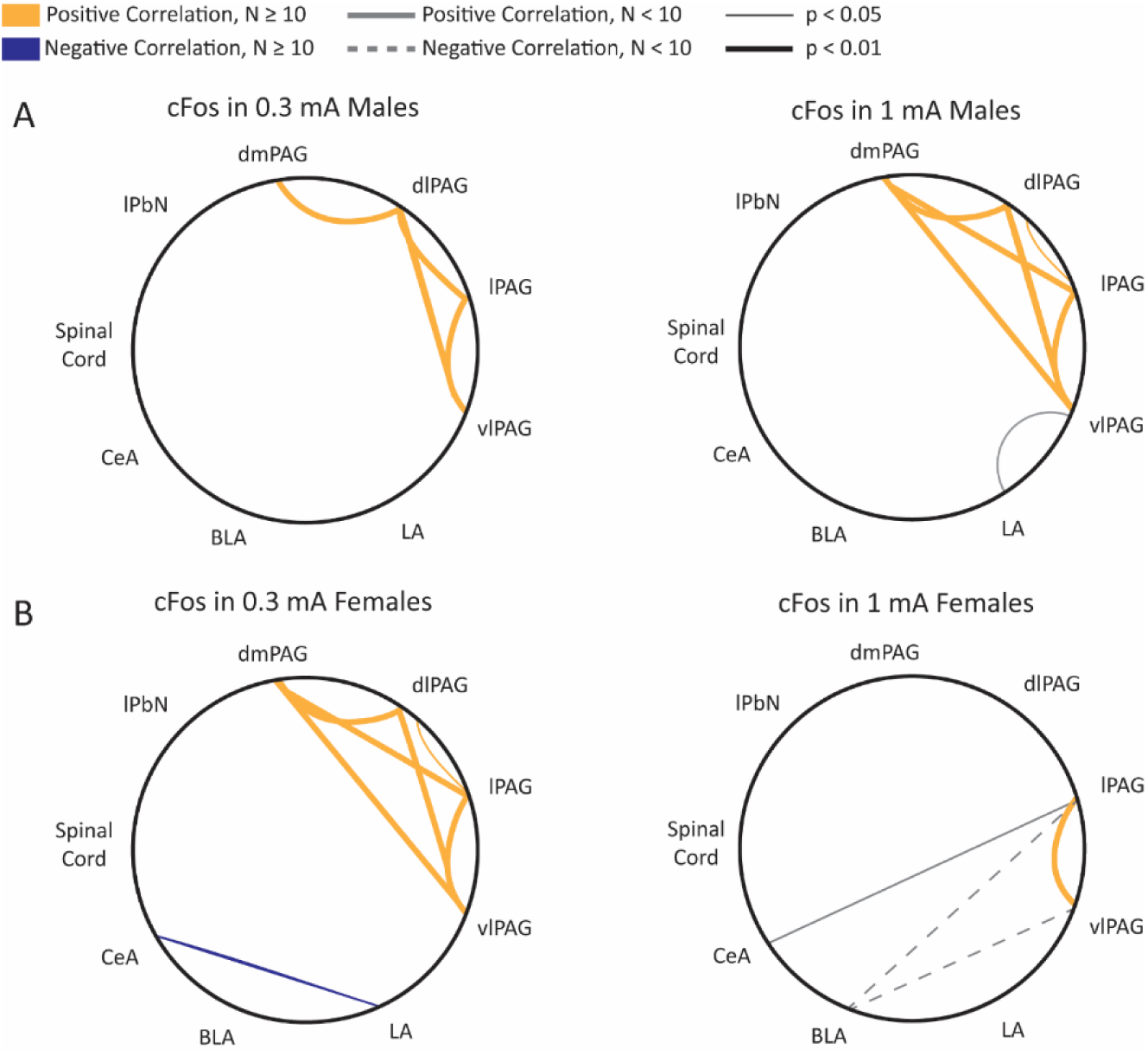
Correlations of cFos expression following fear conditioning of the analyzed regions. **A**. Correlations between males exposed to 0.3 mA shock and 1 mA shock. **B.** Correlations between females exposed to 0.3 mA shock and 1 mA shock. Significant positive correlations shown with yellow connecting lines and significant negative correlations shown with blue connecting line. If N’s for any correlation were less than 10, lines are gray. Solid gray lines indicate positive correlation. Dashed gray lines indicate negative correlation. Level of significance is indicated by the thickness of the line.

## 4. Discussion

Our work strives to broaden understanding of the various types of conditioned fear responses that exist, thereby furthering the inclusion of a wider variety of responses in classical Pavlovian fear conditioning studies. The purpose of these studies was to determine if Darters exhibited increased responding and sensitivity to aversive stimuli, and if this could explain Darters’ heightened unconditioned responses. We compared CORT levels following fear conditioning, behavioral responding in the acoustic startle test, paw withdrawal latency in the hot plate test, and fear conditioning-induced brain activity across Darters, Non-darters, and male rats. Overall, we found that results on these measures do not explain the heightened shock response found in Darters compared to Non-darters and males, with a few notable exceptions.

Experiments 1 and 2 found that Darters do not have higher CORT responses to fear conditioning than Non-Darters, and that their propensity for escape-like movements during fear conditioning does not translate to an increased startle amplitude. In fact, it appears that Darters are slower to startle and have a lower startle amplitude than Non-darters and males. Taken together, these results suggest that Darters are not more sensitive to stress overall, and their choice of escape-like movement during fear conditioning is not due to broad stress or fear-related hypersensitivity.

Experiment 3 furthers our understanding of the relationship between conditioned responses and aversive stimuli and asked if thermal pain was predictive of behavior during fear conditioning. The hot plate test is typically used as a measurement of thermal pain sensitivity in rodents (Espejo & Mir, 1993; Gunn et al., 2011). We replicated previously found results that males have quicker latency to withdraw hind paw than females, (Gunn et al., 2011). Results from the hot plate test also revealed that there might be some differences in sensitivity between fore paw and hind paw in Darters compared to males and Non-darters. It is important to note that the hot plate produces thermal pain, and the foot shock does not, so relationships between behaviors on these assays should be interpreted as exploring individual differences between Darters, Non-darters and males across a range of aversive stimuli, and not as a general indication of Darters’ pain sensitivity. To gain a deeper understanding of the relationship between Darter identity and pain sensitivity, future studies could compare behavior on fear conditioning to behaviors on other commonly used measurements of pain sensitivity, like the Von Frey test examining mechanical sensitivity (Deuis et al., 2017; Minett et al., 2011; Modi et al., 2023), Complete Freund’s Adjuvant for inflammatory pain (Minett et al., 2011; Yu et al., 2021), or in models of chronic pain (Gupta et al., 2017; Kuner & Kuner, 2021; Minett et al., 2011; Raver et al., 2020).

The dorsal horn of the lumbar spine is the primary site of nociceptive input (Coghill et al., 1991; D’Mello & Dickenson, 2008; Wang et al., 2022) and the lack of significant cFos activity based on shock, sex, or Darter identity at the level of the DHL suggests that the fear conditioning paradigm used in this study was not sufficient to activate the canonical peripheral pain pathways beginning at the DHL (Millan, 2002; Westlund & Willis, 2015). This might be due to experimental limitations rather than a true lack of effect. Specifically, the foot shock used in this experiment is 0.5 seconds in duration and each animal is exposed to the shock a total of seven times, which might be too transient of a stimulus to activate cFos expression in the spinal cord. The DHL responds to noxious stimuli (D’Mello & Dickenson, 2008; Watkins et al., 1984), defined as anything potentially damaging or harmful to tissue (Todd, 2010). The shock intensities used in this experiment therefore might not be sufficiently noxious to activate the DHL neurons to protect the limbs from tissue damage. In contrast, the shock intensity effect in the lPbN in females suggests that this region can detect foot shock intensity and that the effect increases as shock intensity increases, but that its activity is not dependent on or responsible for differences seen between female Darters and Non-darters. These results suggest that the DHL is not involved in processing the foot shock, but the primary supraspinal target of nociceptive transmission from the DHL, the lPbN, (Chiang et al., 2019) might be.

The PAG plays a critical role in pain and threat / fear processing. Following fear conditioning, others have shown that each column of the PAG shows an increase in cFos expression (Carrive et al., 1997). Even though the role of the vlPAG in conditioned freezing behavior is thought to be well-established (De Oca et al., 1998; Keay & Bandler, 2015; Liu et al., 2022; Reis et al., 2023; Vianna et al., 2001; Watson et al., 2016), we did not find that 1 mA animals had more cFos expression than 0.3 mA animals, even though our 1 mA animals froze more than our 0.3 mA animals (Figures 3F,G). It is important to note that both 0.3 mA and 1 mA shock intensity groups did have higher levels of cFos than CS-only baseline (dotted line in Figure 6H), so perhaps the vlPAG simply was not able to differentiate between shock intensities. Somewhat surprisingly, the dmPAG and dlPAG showed little to no differences in expression in response to different shock intensities, across sexes, or across Darter identity. The dmPAG and dlPAG are known to be involved in escape-like responses (Deng et al., 2016; Kim etal., 2013; Lefler et al., 2020; Reis et al., 2021; Vianna et al., 2001; Watson et al., 2016), so we expected to see differences between Darter identity groups in these areas. Although, because of the dmPAG’s known role in pain processing (Butler et al., 2011; Linnman et al., 2012), its lack of involvement further suggests that pain from the foot shock is not a motivating driver of the differences seen between males and females and Darters and Non-Darters during fear conditioning. The lack of effect found here might also be due to our choice of slice (–8.04 from bregma) for cFos analysis. The PAG in the rat is known to have different functions depending on the rostral / caudal location (Depaulis et al., 1992; Keay & Bandler, 2015). Regions more rostral or caudal to the bregma point that we chose might have more of an influence. Indeed, many papers do not list their bregma points used for immunohistochemistry procedures (Canteras & Goto, 1999; Comoli et al., 2003; Samineni et al., 2017), or chose sections that were either rostral (De Oca et al., 1998) or caudal (de Mello Rosa et al., 2022) to ours, or included a range of vlPAG coordinates (Borelli et al., 2005). We chose this point because it is close to the anatomical midpoint of the PAG (Loyd & Murphy, 2009), and because of previous studies’ use of this bregma point when investigating pain (Loyd et al., 2007) and fear responses (de Andrade Rufino et al., 2019; Vianna et al., 2001; Wright & McDannald, 2019). The most intriguing and promising column for follow-up studies is the lPAG, in which we observed greater activity in Darters compared to Non-darters. The lPAG is associated with escape and flight behavior, and, in particular, post-encounter and circa-strike defensive behavior, both of which are evasive threat responses (Motta et al., 2017; Zhang et al., 2024). Because of the lPAG’s role in escape behavior in response to threats, it is perhaps unsurprising that we see an effect of conditioned response here, suggesting it may be involved in driving darting as a conditioned response. Higher intensity stimuli have been shown to evoke escape-like responses (Bolles & Fanselow, 1980). It is possible that Darters find the shock to be more aversive or threatening, but not necessarily more painful, which leads to their heightened shock response and escape-like response and therefore heightened cFos expression in the lPAG. Further research into the role of conditioned darting and the lPAG is needed.

Connections between the LA, CeA and BLA are critical for processing both pain and fear and their behavioral expressions (Almeida et al., 2004; Hogri et al., 2022a; Lindsay et al., 2021; Peirs & Seal, 2016). Only the BLA showed a significant effect of shock intensity, highlighting its role in learning the CS-US association and in the processing of aversive stimuli. We would expect more cFos expression following exposure to a 1mA shock because of the increase in shock intensity and resulting increase in freezing, but this was not the case. This suggests that there are other regions responsible for interpreting and communicating the aversive nature of increasing foot shock. In the LA and BLA, there was an increase in cFos expression in the 1 mA female Darters compared to the 1 mA female Non-darters. The LA is involved in the auditory processes involved in formation of the CS-US pairing (Janak & Tye, 2015) and the BLA contains a distinct neural assembly that encodes the negative affective valence of pain (Corder et al., 2019). This supports the hypothesis that Darters might find the foot shock more aversive, leading to a greater recruitment of these areas critical for fear learning and processing the affective side of aversive stimuli. If the stimulus is interpreted as more aversive, Darters might be more motivated to avoid future exposure to it. These results highlight the differential recruitment of brain regions that may lead to similar behavioral outcomes in males and female Non-darters.

Some of the most exciting results came from the differences in our inter-region correlations across the sexes. Males showed similar patterns of correlations of expression, regardless of shock intensity: The columns of the PAG increase in activation together, except for the dmPAG which then increases with the other columns at higher shock intensities. Most of the previous research on function of the PAG has been done in males and, given the known involvement of the PAG in fear conditioning, this broad elevation in expression is unsurprising. Males exposed to a 1 mA shock also show a positive correlation with the LA and vlPAG, which is not seen in the 0.3 mA males, but more N’s would help understand the significance of this correlation.

Females provide a more complex and varied pattern of activation. Females exposed to a 0.3 mA shock show a similar pattern to males, with positive correlations between the sub columns of the PAG, with the addition of a negative correlation between the LA and CeA. The LA and CeA communicate with each other during fear conditioning (Ciocchi et al., 2010; Davis, 1992; Janak & Tye, 2015), so finding this correlation is not surprising. In response to a 1 mA shock, all but one of the intercorrelations between the sub columns of the PAG disappear. Correlations between the vlPAG, a region critical for conditioned freezing, and the lPAG remain. Freezing increases at higher shock intensities, so it is unsurprising that some correlations with the vlPAG stayed consistent across all groups, but the loss of interconnectivity between the other columns of the PAG suggest that females exposed to more aversive stimuli process the experience differently than males and females exposed to less aversive stimuli. This is further supported by the appearance of negative correlations between the BLA and the lPAG and vlPAG in 1 mA females. This suggests that there is a relationship between the activity in these regions and is supported by the decrease in cFos expression seen in 1 mA females in the BLA and CeA (Fig 7F,I). Future studies should investigate the relationships between the sub columns of the PAG with each other, as well as with the lateral, central, and basolateral amygdala.

One aspect of fear conditioning that this study did not examine was fear-conditioned analgesia. Animals who express conditioned freezing are hypothesized to experience fear-conditioned analgesia, characterized by a reduction in pain sensitivity (Fanselow, 1984). Females do not show the same fear-conditioned analgesia as males, with studies either finding less fear-conditioned analgesia or none in females (Llorente-Berzal et al., 2022; Stock et al., 2001). Darters could represent a subset of females in which fear-induced analgesia wholly fails, resulting in the interpretation of the foot shock as more painful and subsequent heightened shock responses. Future studies will examine this possibility.

A potential limitation of this study is the use of cFos as a proxy for neural activity. Although cFos is widely used and accepted as an indicator of neuronal activation (Bullitt, 1990), it is not temporally specific. As a result, we might be unable to detect significant differences across the sexes, Darter identities, and shock intensities. It is possible that some regions are more active earlier in the fear conditioning paradigm than other regions (such as the LA, involved in learning the auditory CS-US association), and that is why we do not see significant effects of shock intensity. Although methods such as fiber photometry would allow more specific mapping of activation, the methods used in this study allowed us to examine a broad range of regions and networks involved in fear conditioning within individual animals. In addition, the use of cFos did not allow us to specify what types of cells are being activated during fear conditioning. Each of the regions examined are heterogenous with respect to cell-type specificity (Chiang et al., 2020; Ge et al., 2022; Keay & Bandler, 2015; McPherson & Ingram, 2022; Swanson & Petrovich, 1998; Watanabe et al., 2017), and we look forward to addressing the question of cell-type specific activation of these regions in future studies.

## 5. Conclusions

This work furthers our understanding about the differences between Darters, Non-darters and males. The overall results from this research allow us to conclude that Darters are not overall simply more sensitive to aversive stimuli, nor that hyperarousal is what leads to the escape-like conditioned responses and heightened unconditioned responses. This adds to our previous body of work, identifying that the sex-biased nature of darting is not dependent on the estrous cycle (Gruene et al., 2015) or weight of the animals (Gruene et al., 2015; Mitchell et al., 2022). This research further emphasizes the importance of looking beyond freezing as a conditioned response (Chu et al., 2024), especially when female subjects are included. It provides novel insights into the regions that are activated and involved in conditioned fear responses, as well as an understanding of how an animal’s conditioned fear responses might or might not be indicative of their behavior in response to other stimuli. Future studies should explicitly investigate other fear responses aside from freezing, as well as the circuits underlying conditioned and unconditioned fear responses.

## Author contributions

JRM, HB, MAL, RMS conceptualized and designed the experiments. JRM, LV, ST, EB, RC, CC, JK, AP, MCP, MAL conducted the experiments. RC, LZ created and optimized cFos protocol. JRM, MAL wrote cFos macro. JRM, ST, VS, AK, MCP, TV, LZ compiled and counted cFos data. JRM, LV, CC, MAL analyzed data. JRM, LV, RMS wrote the manuscript.

## Acknowledgements

We extend our deepest gratitude to Eliza Greiner for assistance in editing this manuscript and with statistical analyses, and to Larry Deng and Esther Ofielu for helping with Experiment 1. Experimental design figures were created using BioRender.

**Appendix A.** Supplementary data tables 1-4 can be found in Appendix A.

