## Appendix A for "Behavioral and neural correlates of diverse conditioned fear responses in male and female rats"

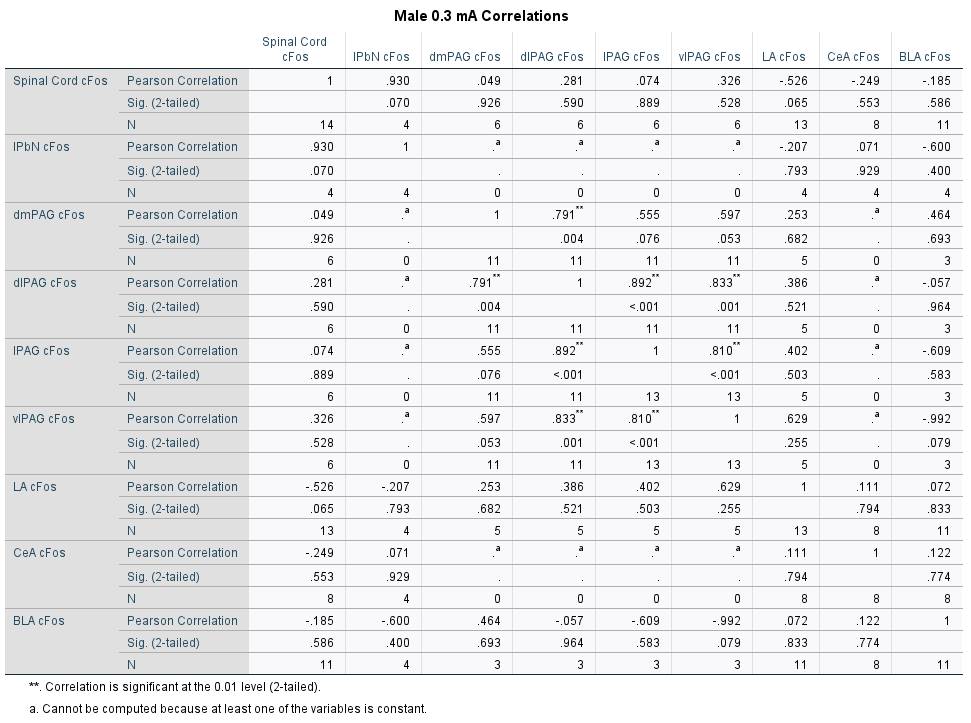

**Table S1. Correlations between all regions examined in 0.3 mA males.**

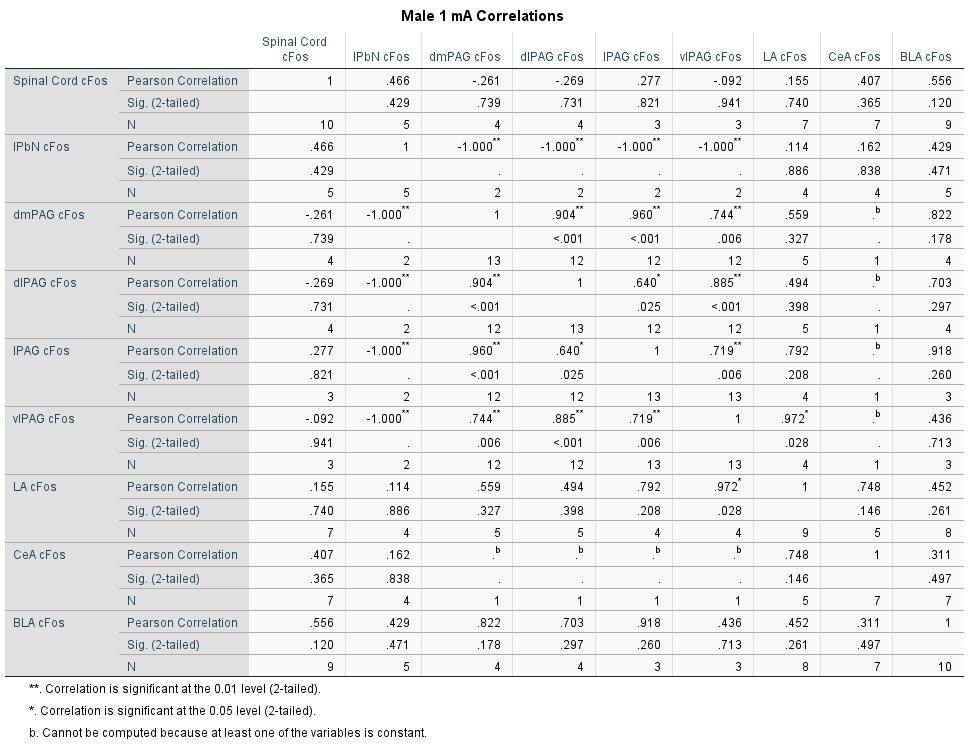

**Table S2. Correlations between all regions examined in 1 mA males.**

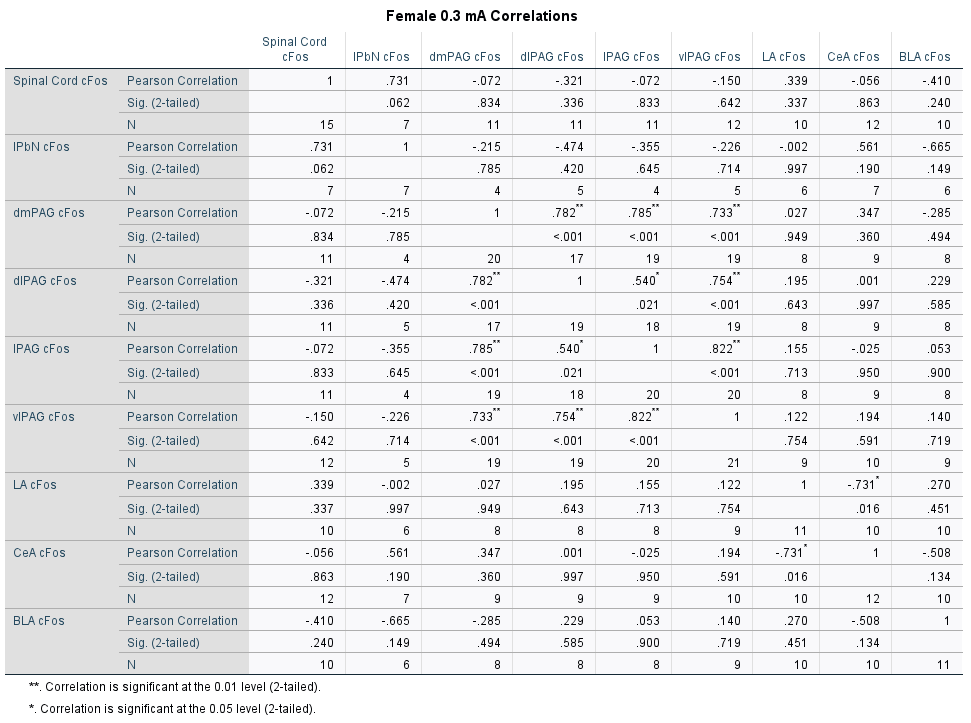

**Table S3. Correlations between all regions examined in 0.3 mA females.**

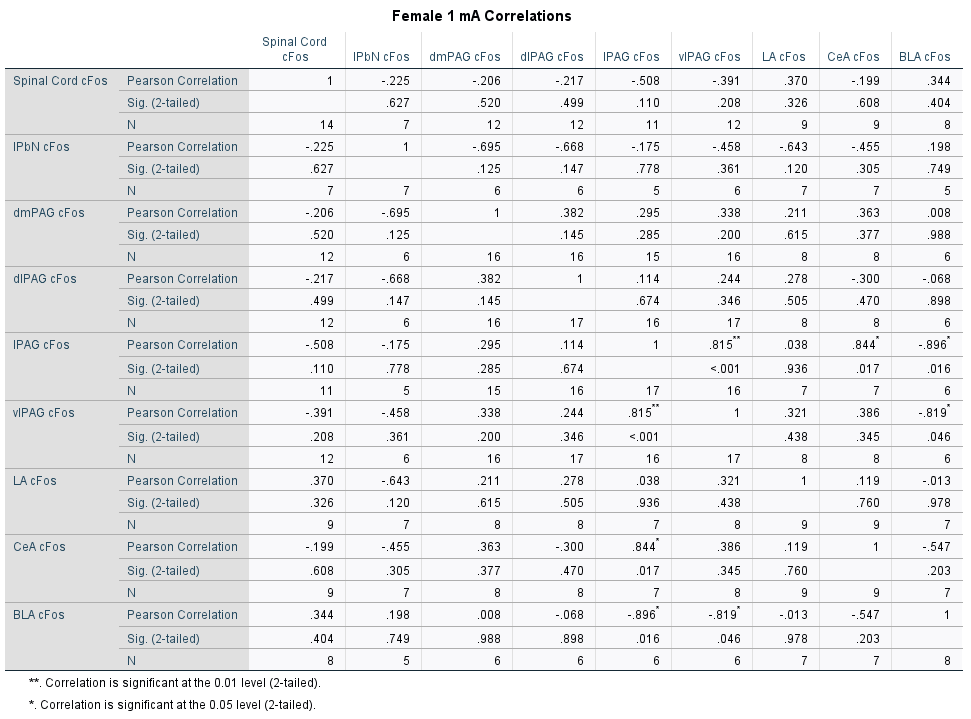

**Table S4. Correlations between all regions examined in 1 mA females.**
